# Synthetic miR-16-5p does not act as a reverse transcriptase co-factor to enhance detection of small RNA

**DOI:** 10.1101/047001

**Authors:** Melissa A. McAlexander, Kenneth W. Witwer

## Abstract

Failure to detect low-abundance microRNAs (miRNAs), for example, in circulating plasma, may occur for a variety of reasons, including presence of enzyme inhibitors. Recently, we received the unusual but intriguing suggestion that miR-16-5p acts as a co-factor of reverse transcriptases, facilitating more efficient reverse transcription of miRNAs and thus enhanced detection of low-abundance miRNAs. We tested this hypothesis by incubating reverse transcriptase with several concentrations of synthetic miR-16-5p and then performing stem-loop RT-qPCR with serial dilutions of miRNA osa-MIR168a. Our results do not support a role for miR-16 as a co-factor of reverse transcriptase.

## INTRODUCTION

Previously, we reported a lack of quantitative PCR (qPCR)-based detection of plant miRNAs, or xenomiRs, in blood of primates that received a plant miRNA-replete dietary substance [1]. Technical controls and digital droplet PCR revealed that any late amplification signal was consistent with non-specific amplification, while synthetic standard curves demonstrated assay sensitivity down to tens or even single digits of copies. These findings were not surprising, since dietary RNA is labile, is exposed to a series of harsh environments in the alimentary canal, and has no well characterized routes of uptake into and distribution within mammals [2-4]. Other groups have reported similar results [5-9] in humans and rodents using a variety of qPCR and sequencing methods.

A colleague who expressed skepticism at our results because of a previous report of xenomiR detection [10] presented an intriguing hypothesis. Specifically, low-level exogenous plant miRNAs in mammals were not detected because the reverse transcriptase we used [acquired from Applied Biosystems by Thermo Fisher (ABI)], required pre-incubation with a small RNA co-factor for optimal activity:

> *“By the way, I guess that you are all using ABI reverse-transcriptase (RTase). If so, I would suggest that (you) mix synthetic mi(R)-16 with RT, incubate for 30m in at 16°C, then use the m ixture to do your qRT-PCR for detecting low level(s) of miRNA (endogenous and exogenous plant miRNA), even low level(s) of mRNA.”*

Although no explanation was given as to why miR-16 might be necessary, we were interested in determining whether this simple additional step prior to reverse transcription and qPCR would lend even more sensitivity to already highly sensitive assays. Therefore, we tested the hypothesis that miR-16-5p acts as a co-factor for reverse transcriptase enzyme.

## MATERIALS AND METHODS

Synthetic oligonucleotides corresponding to hsa-miR-16-5p (sequence UAGCAGCACGUAAAUAUUGGCG, also referred to as “miR-16” or “hsa-miR-16”) and osa-MIR168a-5p (UCGCUUGGUGCAGAUCGGGAC) were ordered from Integrated DNA Technologies (IDT). RT enzyme (MultiScribe^™^ RT enzyme, Applied Biosystems/Life Technologies, now Thermo Fisher) was pre-incubated for 30 minutes at 16°C with synthetic miR-16 at four concentrations: high (400 nM), mid (4 nM), low (400 fM), and none. A dilution series of synthetic osa-MIR168a standard was prepared, with dilutions separated by two-fold (dilutions 1 and 2) and four-fold (dilutions 3-6). Following RT reaction, dilution per manufacturer’s instructions, and preparation of the second-step amplification reactions, there were an estimated 10,000, 5,000, 1,250, 80, and 20 copies of synthetic osa-MIR168a per qPCR reaction. Two-step RT-qPCR assays were from Applied Biosystems/Thermo Fisher under Catalog number # 4427975 (inventoried miRNA assays): hsa-miR-16-5p (000391) and osa-MIR168a-5p (007594_mat). RT and qPCR steps were performed as described previously [11] with a CFX96 qPCR system (BioRad).

## RESULTS

When MIR168a was measured in dilution series, as expected, cycle of quantitation (Cq) values for the first two standard dilutions were separated by approximately one cycle, while Cq for the remaining dilutions were separated by approximately two cycles (Figure 1). The greatest variability was observed at the highest standard dilution, also as expected. At the highest concentration of synthetic miR-16, slightly later amplification (by around half a cycle on average for the first five standard dilutions) was observed. miR-16 was also assayed to confirm the presence of the presumed cofactor (Figure 1).

**Figure 1.**
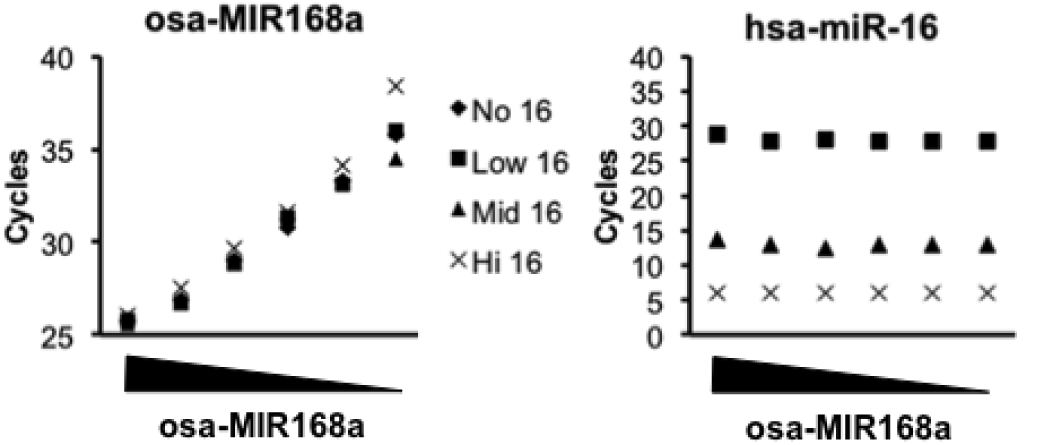
No enhancing effect of miR-16-5p as a reverse transcriptase co-factor. Synthetic miR-16-5p was incubated with reverse transcriptase (Thermo Fisher) for 30 minutes at 16°C at the following concentrations: none (“No 16), 400 fM (“Low 16), 4 nM (Mid 16), and 400 nM (Hi 16). Reverse transcription reactions were performed to detect serially diluted synthetic osa-MIR168a (left) or hsa-miR-16-5p (right) in the same reactions.

## DISCUSSION

These results do not support the hypothesis that synthetic miR-16 acts as a reverse transcriptase co-factor to enhance reverse transcription and subsequent detection of miRNA such as plant miRNA osa-MIR168a in standard stem-loop reverse transcription/hydrolysis probe qPCR assays. Indeed, at the highest pre-incubation concentration of miR-16, there was a decrease in sensitivity. The performance of some TaqMan assays (including, in our hands, the osa-MIR168a and ath-MIR156a assays), may leave only limited room for improvement, and that within a low concentration range that for most source materials would have no relevance for canonical miRNA-mediated regulation: hundreds to thousands of copies per cell [12]. Although one could postulate that miRNAs that we did not test here would be better reverse-transcribed in the presence of miR-16, it is unclear how this effect would be mediated, and we do not plan to investigate this hypothesis further.

## AUTHOR CONTRIBUTIONS

KWW planned the study. MAM performed the experiments. KWW and MAM analyzed the data. KWW wrote the manuscript, and MAM read and approved the manuscript.

## ACKNOWLEDGMENTS

The authors thank members of the Witwer lab for assistance and helpful discussions, and the unnamed colleague for the interesting suggestion of hsa-miR-16-5p as a reverse transcriptase co-factor.

